# Human cytomegalovirus-encoded G protein-coupled receptor (GPCR), UL78, regulates viral reactivation

**DOI:** 10.1101/2025.06.04.657869

**Authors:** Samuel A. Osanyinlusi, Vargab Baruah, Ian J. Groves, Katherine H. Kulp, Benjamin A. Krishna, Christine M. O’Connor

**Author notes:** Address correspondence to Christine M. O’Connor. Current address: Department of Medicine, University of Cambridge, Cambridge CB2 0QQ, UK.

## Abstract

Human cytomegalovirus (CMV) is a ubiquitous pathogen that establishes life-long, latent infection in hematopoietic cells. Immune-competent individuals are usually asymptomatic for disease. However, immune dysregulation in latently-infected individuals can result in viral reactivation, often causing further complications. Viral gene transcription during latency is restricted, although the CMV-encoded G-protein coupled receptor homologs, US28 and UL78, are expressed. We and others find US28 is critical for establishing and maintaining viral latency, in part, through regulating host cell signaling. How US28 switches from pro-latent to pro-lytic during reactivation, however, is unknown, though our findings herein reveal a role for UL78. Myeloid cells infected with a UL78 ORF deletion mutant maintain viral latency yet fail to efficiently reactivate. However, the UL78 G protein-coupling domain is not required for reactivation, suggesting UL78-mediated signaling is not critical for reactivation. Prior work revealed UL78 and US28 interact, resulting in altered US28-mediated signaling. Additionally, we showed US28 attenuates ERK phosphorylation during latency, while ERK is phosphorylated upon reactivation; however, the mechanism underlying this switch is unknown. Thus, we hypothesized the UL78:US28 interaction is important for altering US28-mediating signaling upon viral reactivation. We find US28 and UL78 interact during lytic infection of fibroblasts and colocalize in myeloid cells upon their differentiation. Further, reactivation in myeloid cells latently-infected with wild-type virus results in upregulated ERK phosphorylation, while parallel cultures infected with the UL78-deficient virus fail to do so. Our data reveal the first function for UL78 in myeloid cells, where it influences cellular signaling to switch from pro-latent to pro-lytic.

**Importance:** Cytomegalovirus (CMV) is a ubiquitous human herpesvirus, infecting the majority of the population worldwide. As with all herpesviruses, once an individual is infected with CMV, the virus remains in a person’s blood cells for their life in a silenced state called latency, and this infection, for the most part, remains asymptomatic. When an infected individual’s immune system fails to function properly, however, CMV can become active (termed viral reactivation), which allows the virus to replicate and cause downstream disease. Our understanding of the cellular and viral factors that dictate this switch from silenced to activated remains incomplete. Here, we show a viral protein, UL78, is required for this switch. We find UL78 helps to reshape cellular signaling, changing the cell environment from one that favors latency to one that instead supports reactivation. This highlights a new avenue for therapeutic intervention to prevent CMV reactivation and downstream disease.

## INTRODUCTION

Human cytomegalovirus (CMV) is a ubiquitous, betaherpesvirus pathogen with a high global prevalence. Seroprevalence ranges from 50% to over 90%, depending on socio-economic factors and geographic region (1, 2), and thus, CMV remains a major public health threat. Like all herpesviruses, CMV infection is life-long, characterized by its ability to establish latency in host cells, with periodic reactivation upon immune dysregulation. Primary infection of healthy individuals is usually self-resolving with little or no symptoms; however, CMV reactivation in immunocompromised (e.g., AIDS) and immunosuppressed (e.g., organ transplant recipients) individuals is characterized by severe disease (3). Additionally, congenital CMV infection in the immunonaïve is a leading cause of birth defects globally, with accompanying developmental disabilities, such as microcephaly, hearing loss, and cognitive impairment (4).

CMV establishes latent infection in cells of the hematopoietic lineage, including bone marrow-derived CD34^+^ hematopoietic progenitor cells (HPCs) and circulating CD14^+^ monocytes (5). During the maintenance of latency, the viral major immediate early (MIE) locus, which includes the MIE promoter (MIEP) and alternative promoters that drive the IE genes (6–9), *UL123* and *UL122*, is largely repressed (10). Repression of this locus, and ultimately the transcription of these IE genes and their translated proteins, is critical to maintaining CMV genomic silencing required to maintain latency. However, cellular differentiation triggers cues that alter the host cell environment, making it more amenable to viral reactivation. Such changes impact the MIE locus, switching it from a repressed enhancer/promoter region to one that is more active. This is achieved through chromatin modifications and chromatin-associated proteins/factors, three-dimensional reorganization of the viral genome, and changes to cellular signaling, which influences the recruitment of transcription factors that regulate the activity of the MIE enhancer/promoter locus. The switch between latency and reactivation is thus hinged on the regulation of the MIE region, coordinated by an intricate interplay between cellular and viral factors, as well as host cell signaling. However, a complete understanding of the virus- and host-encoded proteins that regulate this balance remains incomplete.

CMV has the capacity to encode over 200 open reading frames (ORFs), though many of these genes display reduced transcriptional profiles during latent infection. CMV has no distinct latency transcriptional profile (11–14), although there are certain genes that retain increased expression over others during latency, including three of the four CMV-encoded G protein-coupled receptors (GPCRs): US28, UL33, and UL78 (the fourth viral GPCR [vGPCR], US27, is repressed during latency (14, 15)). Like host-encoded GPCRs, the vGPCRs have been studied for their signaling capabilities and how their signaling reshapes the cellular milieu. More recently, our lab and others have begun to investigate how the latently-expressed vGPCRs function during CMV latency and/or reactivation. We and others showed US28 is critical for establishing and maintaining latency (15–25), while UL33 is important for efficient viral reactivation (26), and in each case, these vGPCRs alter host cell signaling. Unlike US28 and UL33, however, investigators have not yet found signaling properties that are attributed to UL78. In line with this, UL78 is also an orphan receptor, as no known ligands bind this vGPCR to potentiate signals. Further, UL78 has distinctive structural peculiarities, including a long C-terminal tail and a ‘DRL’ motif (instead of the canonical DRY) as its predicted G protein-coupling domain, which may prevent UL78 from coupling G proteins (27). Despite this, UL78 is important for efficient viral entry and in turn, replication in epithelial cells (28). Similarly, the rodent CMV orthologues, M78 and R78, encoded by murine CMV and rat CMV, respectively, are also critical for efficient pathogenesis in their hosts (29–32). Thus, while UL78 may not directly potentiate cell signaling, its expression is clearly important for infection. Further, UL78 may impart function through interactions with other viral proteins. Indeed, ectopic co-expression of UL78 and US28 results in their heteromerization, leading to altered US28-mediated signaling (33), though whether this occurs in the context of infection remains outstanding. As *UL78* transcripts are detected during natural and experimental latency (11, 13), we reasoned that UL78 may function during this phase of infection, possibly interacting with other vGPCRs, such as US28, in turn impacting downstream US28-mediated signaling.

Herein we show UL78 is critical for efficient reactivation of CMV from latency. Our data reveal UL78 protein is expressed in *in vitro* (THP-1 cells) and *ex vivo* (primary CD14^+^ monocytes) models of CMV latency. Further, our data show a complete UL78 ORF deletion virus fails to efficiently reactivate from latency despite maintaining viral genomes; however, this deficiency is not dependent upon UL78’s DRL G protein-coupling motif, as mutation of this motif does not deter efficient viral reactivation. We also find UL78 and US28 colocalize upon myeloid cell differentiation/viral reactivation, as well as interact in lytically-infected fibroblasts. Consistent with prior work (33), ectopic co-expression of these two vGPCRs in fibroblasts results in altered US28-driven signaling. Importantly, UL78 expression impacts US28-mediated signaling upon reactivation in myeloid cells. While reactivation of wild type virus results in the upregulation of ERK phosphorylation, consistent with our prior work (22, 25), reactivation of a UL78 ORF deletion virus fails to result in robust ERK phosphorylation. Collectively, our findings suggest UL78 expression in differentiated myeloid cells influences cellular signaling to switch from pro-latent to pro-lytic.

## RESULTS

### UL78 is expressed during CMV latency and is required for efficient viral reactivation

Prior work revealed *UL78* transcript is expressed during both experimental and natural latency (11, 13), though whether the protein is expressed and if it functions during latency and/or reactivation remained outstanding. We first evaluated the abundance of UL78 protein during latency in infected THP-1 and primary CD14^+^ cells. To this end, we infected cells with TB40/E*mCherry* (wild type; WT) or TB40/E*mCherry*-UL78-3xF (UL78-3xF) under latent conditions for 7 days (d). We also included lysates from UL78-3xF-infected fibroblasts (NuFF-1) as a positive control. In parallel, to ensure our cultures were latent, we probed for pp65, as this protein is abundant during lytic infection and repressed during latency. Our data reveal UL78 is expressed during latent infection in THP-1 (**Fig. 1A**) and CD14^+^ (**Fig. 1B**) cells in the absence of pp65, though UL78 is less abundant than during lytic expression of this protein (**Fig. 1**).

**Figure 1.**
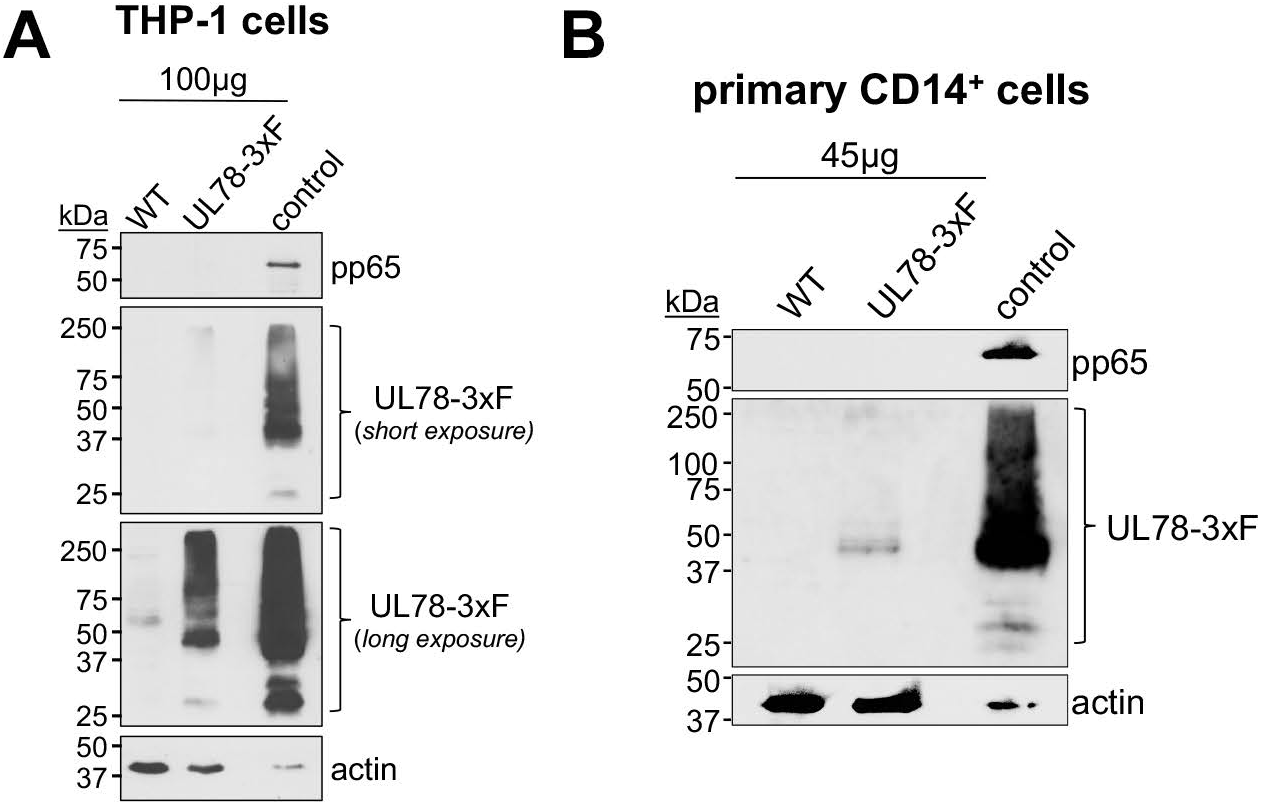
UL78 is expressed during latent infection of myeloid cells. **(A)** THP-1 or **(B)** primary CD14^+^ cells were infected with the indicated viruses (MOI = 1.0 TCID_50_/cell) for 7 d under latent conditions. **(A, B)** Whole cell lysates were collected and probed with antibodies directed at pp65, FLAG (to detect UL78), or actin. Control lysates from UL78-3xF-infected NuFF-1 cells are shown in the right lane (20µg) of each blot as a control. Representative blots, **(A)** n = 3; **(B)** n = 2.

We next tested the requirement for UL78 expression during latency and/or reactivation. To this end, we leveraged our previously characterized UL78 ORF deletion virus, TB40/E*mCherry*-UL78Δ (UL78Δ) (28). Since UL78-mediated signaling has not been evaluated in myeloid cells, we generated three additional UL78 mutant viruses wherein we mutated the putative DRL G protein-coupling domain using BAC recombineering (34, 35): 1) TB40/E*mCherry*-UL78^AAA^-3xF (UL78^AAA^), mutating DRL to AAA; 2) TB40/E*mCherry*-UL78^DAL^-3xF (UL78^DAL^), mutating DRL to DAL; and 3) TB40/E*mCherry*-UL78^DRY^-3xF (UL78^DRY^), mutating DRL to the more canonical DRY motif. In line with our prior work (28), UL78Δ displayed wild type lytic growth relative to WT and UL78-3xF, as did our newly generated UL78 mutants (**Fig. S1A**). We also confirmed the newly generated UL78 mutant viruses expresses the UL78 protein, as measured by FLAG expression (**Fig. S1B**). To evaluate the function of UL78 during latency and/or reactivation, we infected primary CD14^+^ monocytes with our panel of viruses for 7 d under latent conditions, after which half of the cultures were maintained under latent conditions, while the remaining half were treated with M-CSF to differentiate the cells and reactivate the virus. We then co-cultured infected CD14^+^ cells with fibroblasts for a further 14 d to quantify the frequency of infectious centers by extreme limiting dilution analysis (ELDA) (36). As expected, UL78-3xF displayed an increase in the frequency of infectious centers indistinguishable from WT infected cells treated with M-CSF. (**Figs. 2, S2**; blue vs. gray checkered bars). However, UL78Δ-infected CD14^+^ cells treated with M-CSF fail to show a robust increase in infectious center frequency compared to non-treated cultures. Compared to WT- and UL78-3xF-infected cells treated with M-CSF, UL78Δ-infected cells show a less than 2-fold increase in viral reactivation (**Figs. 2, S2**; green vs gray and blue checkered bars). Finally, UL78^AAA^-, UL78^DAL^-, and UL78^DRY^-infected cultures treated with M-CSF resulted in a significant increase in the frequency of infectious centers relative to parallel cultures in the absence of this cytokine (**Figs. 2, S2**). While there is statistical significance between UL78-3xF- and UL78^AAA^-infected cultures treated with M-CSF (**Fig. 2A**), it is clear this virus efficiently reactivates from latency. Further, UL78^DAL^- and UL78^DRY^-infected cultures reactivate to infectious center frequencies that are indistinguishable from either WT or UL78-3xF (**Fig. 2B**). These data suggest the putative G protein-coupling domain is not required for efficient viral reactivation. This is consistent with our current understanding of this vGPCR, as there are no studies to-date that show signaling properties for UL78 (27, 37).

**Figure 2.**
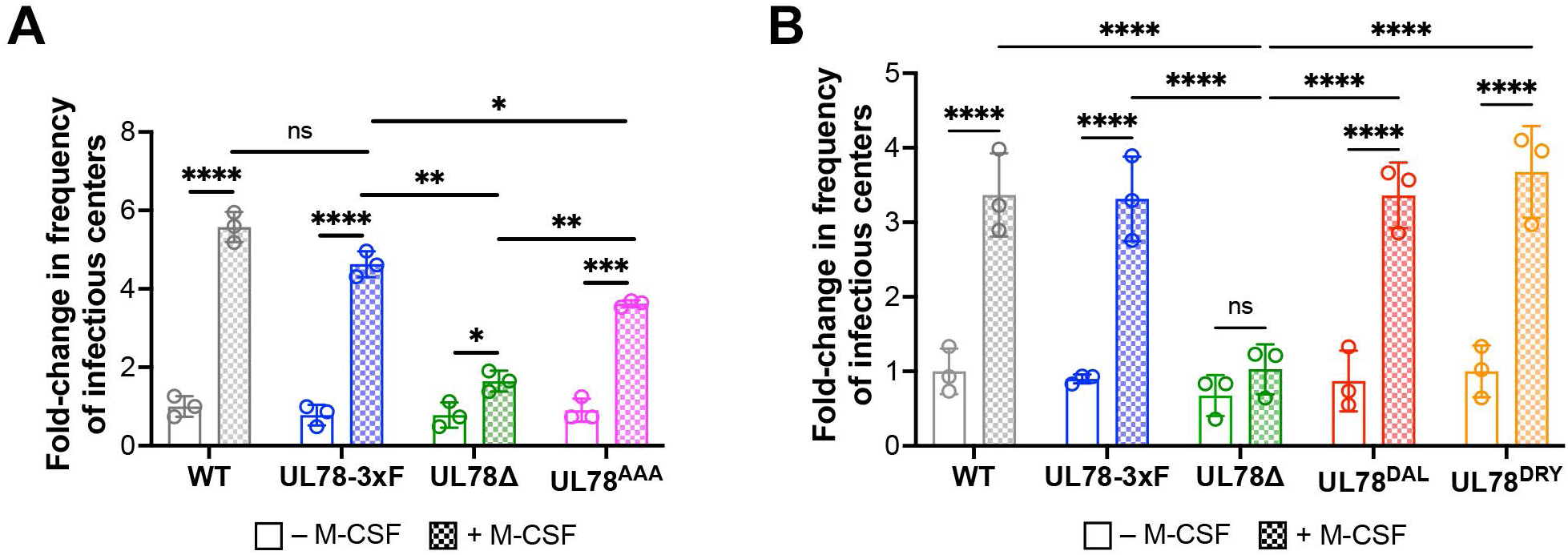
UL78 is important for efficient viral reactivation, independent of its putative G protein-coupling domain. Primary CD14^+^ cells were infected (MOI=1 TCID_50_/cell) with **(A)** WT, UL78-3xF, UL78Δ, or UL78^AAA^ or **(B)** WT, UL78-3xF, UL78Δ, UL78^DAL^, or UL78^DRY^. **(A,B)** Infected cells were then cultured for 7 d in conditions favoring latency. CD14^+^ cells were maintained under latent conditions (-M-CSF) or treated with M-CSF (+ M-CSF) to differentiate the cells and then co-cultured with naïve fibroblasts. Following 14 d in co-culture, ELDA was used to quantify reactivation. Data is shown as fold-change in the frequency of infectious centers, relative to WT latent cultures (open gray bar). Data points (open circles) represent the mean of a biological replicate, each with three technical replicates (see Supplementary Figure 2); error is shown as SD of the mean of the three biological replicates. Statistical significance calculated using T-test; data is not significant unless indicated. * *p<*0.05, ** *p<*0.01, *** *p<*0.0005, **** *p<*0.0001

The failure of UL78Δ-infected cells to reactivate suggests UL78 is required for efficient maintenance of latency or reactivation from latency. Thus, to delineate these possibilities, we evaluated the ability of each virus to maintain viral latency in CD14^+^ cells. To this end, we infected cells under latent conditions with WT, UL78-3xF, or UL78Δ, and assessed viral genomes at 2 and 7 dpi. Our data reveal UL78Δ-infected cells maintain genomes comparable to both WT- and UL78-3xF-infected CD14^+^ cells (**Fig. 3**). Collectively, these data indicate UL78 expression is required for efficient viral reactivation in CD14^+^ cells.

**Figure 3.**
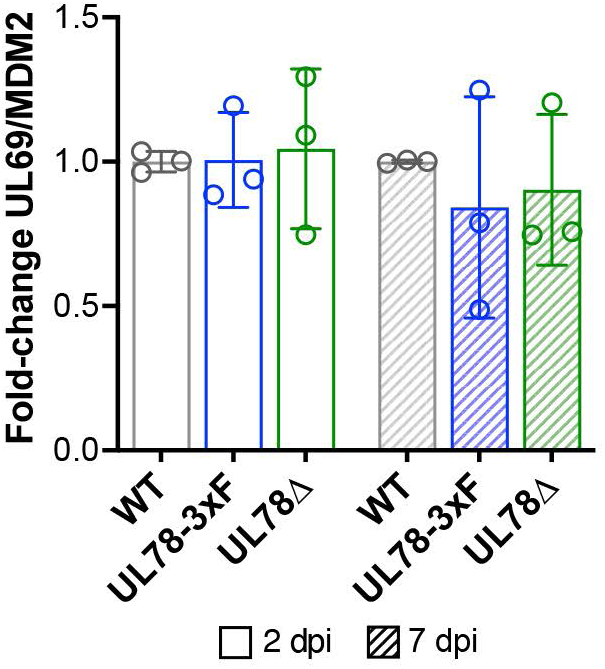
UL78 does not impact genome maintenance in hematopoietic cells. Primary CD14^+^ cells were infected with WT, UL78-3xF, or UL78Δ (MOI = 1 TCID_50_/cell) and cultured for 2 (open bars) or 7 (slanted bars) days under latent conditions. Cells were then harvested, DNA isolated, and viral and cellular genome copies were quantified by qPCR using primers directed at the CMV UL69 non-promoter region and cellular MDM2, respectively. The ratio of UL69 to MDM2 is plotted as arbitrary units (AU) and is depicted as fold-change relative to WT (gray) at each time point. Data points represent mean values from three biological replicates, each with three technical replicates. Error bars denote SD of three biological replicates. Statistical significance was calculated by two-way ANOVA with Tukey’s multiple comparisons test. Data is not statistically significant.

### UL78 and US28 proteins interact and colocalize during CMV infection

Since expression of UL78 is important for efficient viral reactivation, we next asked how this vGPCR influences the switch from latent to lytic infection. Tschische, et al. demonstrated previously that ectopic expression of US28 and UL78 together in HEK293T cells resulted in their heteromerization and in turn, altered US28-mediated cell signaling (33). We and others have shown US28-potentiated signaling is critical to maintain latency (17, 18, 22, 24, 25), yet how these signals are tempered or altered to support viral reactivation is elusive. Thus, we hypothesized that UL78 functions to interact with US28 during viral reactivation, in turn altering pro-latent US28-mediated signaling to that which is more supportive of reactivation/lytic infection. To begin to interrogate this possibility, we first evaluated the potential interaction between US28 and UL78 in lytically infected fibroblasts. To this end, we generated a dual-tagged virus, TB40/E*mCherry*-US28-3xF:UL78-3xHA (US28-3xF:UL783xHA) using the TB40/E*mCherry*-US28-3xF (US28-3xF) backbone, wherein we engineered an HA triple epitope tag at the C-terminus of UL78. This newly generated virus displays wild type lytic growth compared to WT and US28-3xF (**Fig. S3A**) and expresses each viral protein, which we assessed using antibodies to the respective epitope tags (**Fig. S3B**). We then used this virus to test the interaction of US28 and UL78 during lytic infection. To this end, we infected NuFF-1 fibroblasts with US28-3xF:UL78-3xHA and confirmed the interaction of these two vGPCRs by co-immunoprecipitation (**Figs. 4, S4**). While our data reveal US28 and UL78 interact in the context of lytic infection, this does not necessarily mean this interaction occurs during myeloid cell infection. US28 (15) and UL78 (**Fig. 1**) are both expressed in myeloid cells, although the level of their expression is low, requiring significant protein concentrations for detection. Thus, it is challenging to obtain the abundance of myeloid cells to assess the US28:UL78 interaction. Therefore, we instead evaluated the colocalization of these proteins by immunofluorescence assay (IFA) in myeloid cells using two, parallel approaches. First, we infected THP-1 cells with WT or US28-3xF:UL78-3xHA under latent conditions for 7 d, and then differentiated the cells with TPA for an additional 2 d. Second, we pre-treated THP-1 cells with TPA to differentiate the cells for 1 d, after which we infected the cultures with either WT or US28-3xF:UL78-3xHA for an additional 2 d. In each case, cells were then fixed, permeabilized, and stained with antibodies directed at FLAG (US28) and HA (UL78). Upon cellular differentiation of latently-infected cells, we observed diffuse cellular staining of both vGPCRs, along with distinct perinuclear colocalization of both UL78 and US28 (**Fig. 5A**), which is consistent with the location of the viral assembly compartment (VAC) (38). We quantified the colocalization of UL78 and US28 using the pixel intensity correlation over space method based on Pearson’s correlation coefficient and found strong correlation of 0.73±0.08 (**Fig. 5C**), suggesting these two proteins colocalize following reactivation stimulus. In cells that were pre-differentiated prior to infection, we also observe colocalization of US28 and UL78, and again note the perinuclear VAC staining, as well as additional, diffuse cellular staining (**Fig. 5B**). US28 and UL78 also colocalize in the VAC in lytically-infected fibroblasts (**Fig. S5**), consistent with our prior work (21, 28). We also quantified the colocalization of US28 and UL78 in pre-differentiated, infected cells, which revealed a Pearson’s R value of 0.69±0.03 (**Fig. 5C**), within the range of moderate correlation (0.4-0.7). It is important to note infection of the cells in **Fig. 5B** were not synched, thus there are inevitably some cells that are further along in the infection cycle than others, as noted by the intensity of the mCherry staining (**Fig. 5B**). Based on our prior work, UL78 is more diffuse at earlier time points, with more focused localization to the assembly complex as the infection progresses (28). We observe the colocalization of US28 and UL78 in the mCherry bright cells (**Fig. 5B**), suggesting cells further along in infection support US28:UL78 colocalization following *de novo* infection of pre-differentiated cells. Collectively, these data suggest US28 and UL78 interact during lytic infection of fibroblasts and colocalize in differentiated myeloid cells that support viral reactivation.

**Figure 4.**
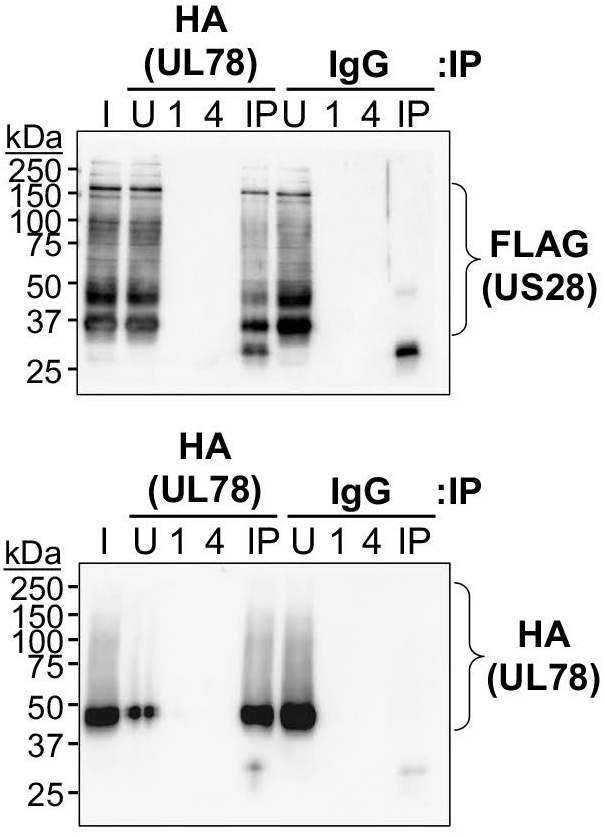
UL78 and US28 interact in lytically-infected fibroblasts. NuFF-1 fibroblasts were infected with TB40/E*mCherry*-US28-3XF:UL78-3xHA (MOI = 0.5 TCID_50_/cell). Cells were harvested 96 hpi and lysates were immunoprecipitated (IP) with either anti-HA or anti-IgG antibodies and immunoblotted for (**top**) FLAG to detect US28 or (**bottom**) HA to detect UL78. I, input; U, unbound; 1, 1^st^ wash; 4, 4^th^ wash; IP, immunoprecipitated sample. Representative blots shown. Control co-IPs from TB40/E*mCherry*-infected cells are shown in Supplementary Figure 4.

**Figure 5.**
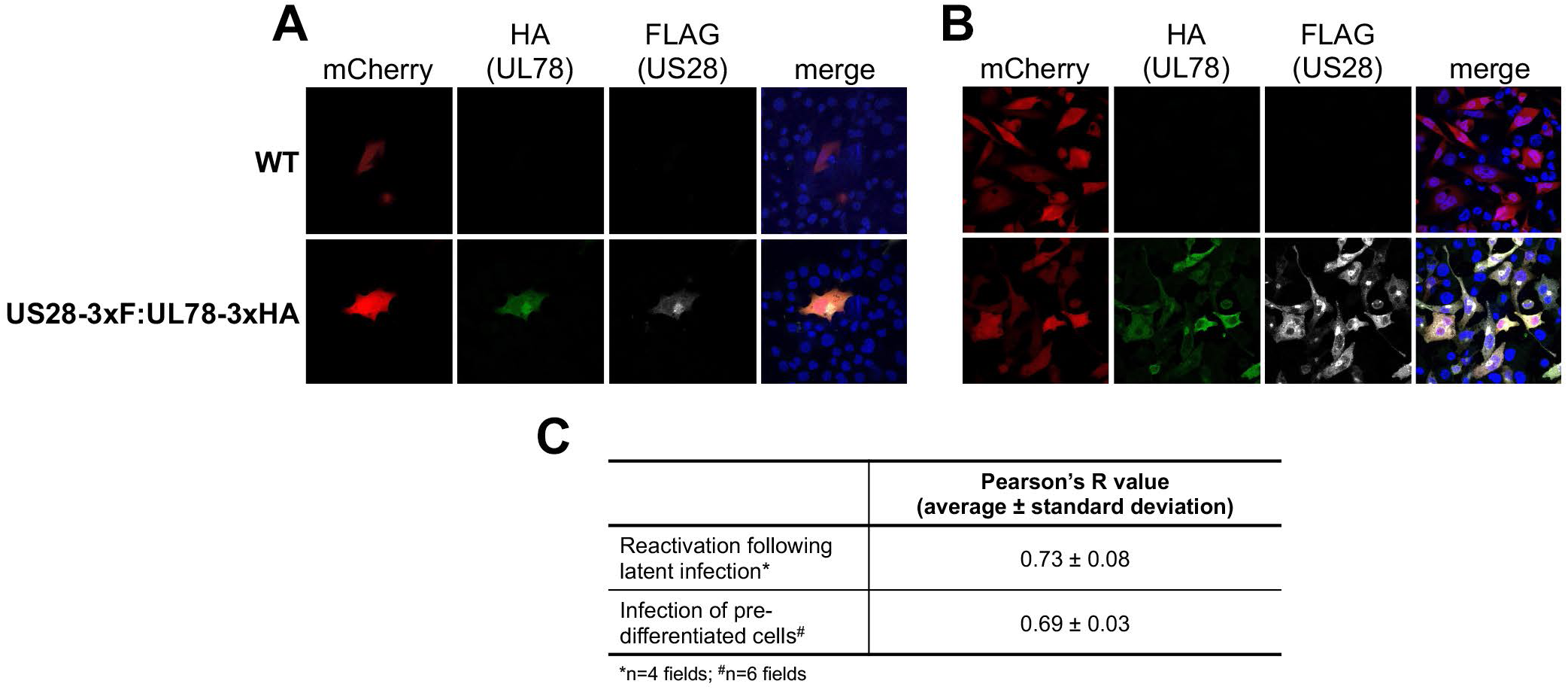
UL78 and US28 co-localize in differentiated THP-1 cells. (**A**) THP-1 cells were infected (MOI = 1.0 TCID_50_/cell) with the indicated viruses under latent conditions for 7 d, after which cells were treated with TPA for an additional 2 d to differentiate the cells. (**B**) THP-1 cells were differentiated for 1 d with TPA, after which cells were infected (MOI = 1.0 TCID_50_/cell) with the indicated viruses for an additional 2 d. (**A, B**) Cells were fixed, permeabilized, and stained with DAPI (to detect nuclei) or antibodies directed at HA (to detect UL78) or FLAG (to detect US28). mCherry is shown as a marker of infection. Images were captured using a 63x objective. N = 3; representative images shown. (**C**) Pearson’s R value was used to calculate the colocalization of US28 and UL78 using images from a representative experiment (0.1-0.3, weak correlation; 0.4-0.7, moderate correlation; >0.7, strong positive correlation).

### UL78 protein expression alters US28-mediated signaling

As mentioned above, heteromerization of US28 and UL78 result in altered US28-mediated signaling in HEK293T cells (33). Thus, based on our data revealing the US28:UL78 interaction in fibroblasts (**Figs. 4, S4**) and colocalization of these vGPCRs in myeloid cells (**Fig. 5**), we next asked if UL78 expression altered US28-mediated signaling. We and others have shown US28 attenuates MAPK signaling during viral latency (18, 22, 25). Thus, we tested whether co-expression of UL78 with US28 altered US28’s ability to repress phosphorylation of ERK, a downstream protein in the MAPK signaling cascade. We transfected fibroblasts with an expression construct for US28-3xF, alone or in combination with one expressing UL78-V5, and assessed ERK phosphorylation. In the presence of UL78, phosphorylation is significantly increased (**Fig. S6**), suggesting co-expression of UL78 with US28 alters US28-regulated ERK attenuation. We also evaluated this phenotype in the context of myeloid cell infection. To this end, we mock-, WT-, or UL78Δ-infected THP-1 cells under latent conditions for 7 d, after which we treated half of the cells with TPA to induce differentiation/reactivation and the other half with DMSO to retain latent culture conditions for an additional 2 d. We then evaluated the phosphorylation level of ERK across the various conditions. Consistent with our previous work and that of others (22, 25, 39, 40), WT-infected myeloid cells display increase in phospho-ERK upon latent infection. Additionally, UL78Δ-infected cells cultured under latent conditions have no statistically significant change in phospho-ERK compared to WT-infected counterpart cultures (**Fig. 6**, -TPA conditions). As expected, upon differentiation of the cells with TPA, we observe a robust increase in phospho-ERK in WT-infected cells (**Fig. 6**). This consistent with previous findings that show ERK phosphorylation is critical for viral reactivation (22, 25, 39–41). This is also similar to US28Δ-infected cells that fail to maintain latency, as these cultures also have increased phospho-ERK, as US28 tempers ERK phosphorylation to a level below a certain threshold to maintain latency (18, 22, 25). In line with this, TPA-treatment of UL78Δ-infected cells did not result in an increase in phosphorylated ERK (**Fig. 6**), suggesting UL78 expression is important for upregulation of ERK phosphorylation, which is otherwise attenuated by US28 during viral latency (18, 22, 25). Collectively, our data reveal UL78 influences cellular signaling to switch from pro-latent to pro-lytic.

**Figure 6.**
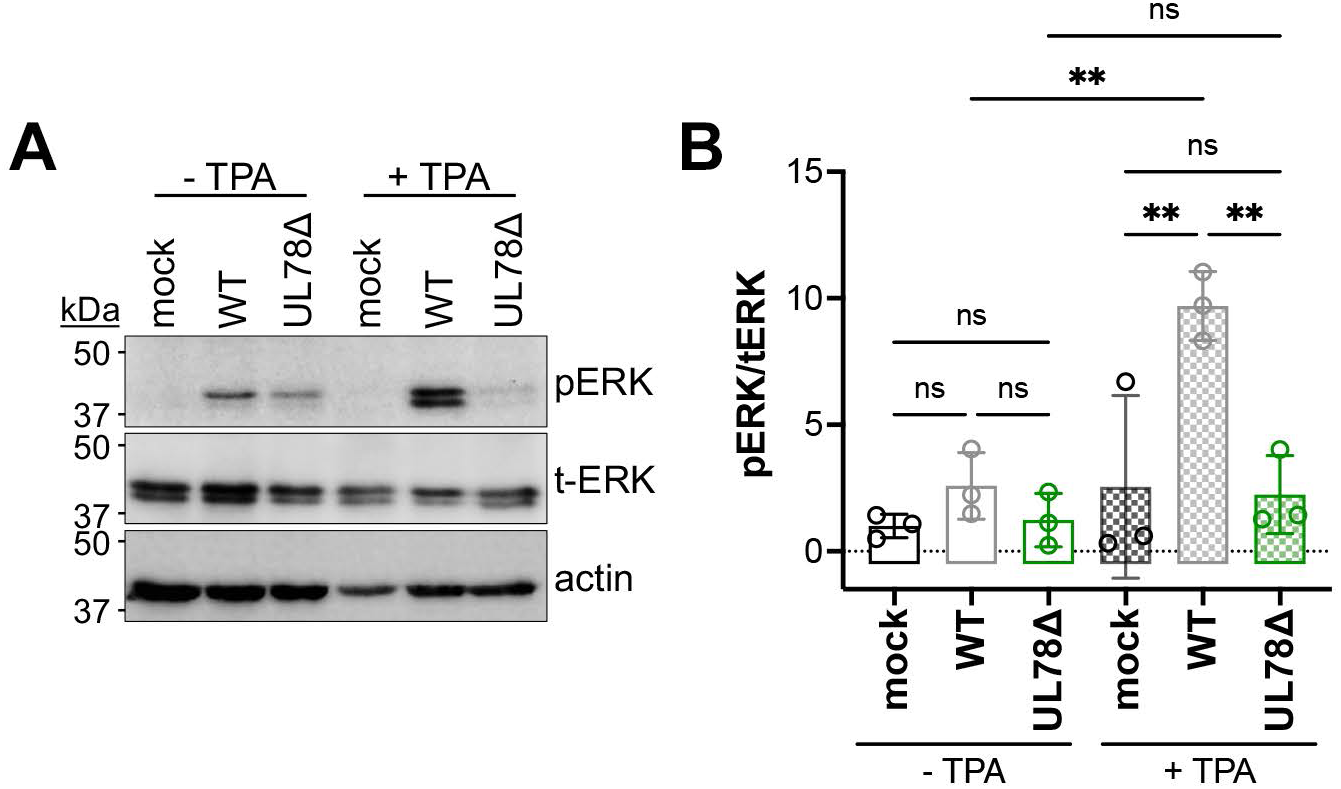
UL78Δ-infected THP-1 cells fail to upregulate ERK signaling upon reactivation. (**A**) THP-1 monocytic cells were mock-, WT-, or UL78Δ-infected (MOI = 1.0 TCID_50_/cell) under latent conditions for 7 d. Cultures were treated with TPA (+TPA) or DMSO (-TPA) for an additional 2 d. Cell lysates were probed with antibodies directed at phosphorylated ERK (p-ERK), total ERK (t-ERK), or actin. N = 3; representative blots shown. (**B**) Data from (A) were analyzed by densitometry to determine the phospho-ratio of pERK/t-ERK. Data points represent the value from each of the 3 biological replicates. Data is shown relative to pERK/t-ERK for control cells (mock, -TPA; set to 1.0). Statistical significance was calculated by one-way ANOVA; error bars represent SD.

## DISCUSSION

Herein, we describe the first function for UL78 in myeloid cells, where this vGPCR influences cell signaling to switch from pro-latent to pro-lytic. We find UL78 protein is expressed in latently-infected THP-1 and primary CD14^+^ cells (**Fig. 1**), where its expression is important for efficient viral reactivation (**Fig. 2**), though is not required to maintain viral latency (**Fig. 3**). Further, we find the UL78 putative G protein-coupling domain is not required for reactivation (**Fig. 2**), suggesting UL78-mediated signaling does not contribute to this phenotype. We also show UL78 interacts with US28 in infected fibroblasts (**Fig. 4**), and these two vGPCRs colocalize in differentiated myeloid cells (**Fig. 5**). Finally, upon reactivation, WT-infected myeloid cells display an upregulation of phosphorylated ERK, while cells infected with the UL78-deficient virus fail to display this increase (**Fig. 6**). Overall, our data reveal a function for UL78 in reshaping the cellular environment from pro-latent to pro-lytic during CMV reactivation.

UL78’s influence on cell signaling supports a mechanism by which the cell, at least in part, shifts from favoring latency to favoring lytic replication. Though we were unable to demonstrate a direct interaction between US28 and UL78 during viral reactivation in myeloid cells, their colocalization in differentiated myeloid cells suggests that they are at least in close proximity, if not physically interacting. This colocalization is particularly evident in cells that were stimulated with TPA following latent infection (**Fig. 5A,C**). Future work will include understanding to which membranes and organelles these vGPCRs colocalize using high resolution microscopy. Also, there are several possibilities for why we were unable to co-IP these two vGPCRs. First, the protein quantities required to detect the interaction could be beyond our capabilities with the latency systems we have. Indeed, detection of UL78 (**Fig. 1**) and US28 (15) require significant protein concentrations, and thus, increased starting material. Compensating for this by increasing concentrations can have deleterious impacts on antibody binding and/or column capturing. Finally, it is also possible these two proteins are simply within close proximity, but do not physically interact, and their abundance and spatial distribution impact the cross-talk. This is not unprecedented, as this indeed occurs with other receptor proteins (e.g., ref. (42)). Nonetheless, we are actively investigating these possibilities as we continue to dissect the impact of UL78 on US28.

It is evident that UL78’s expression is required to alter ERK activity from its US28-driven attenuated form during latency, to one that has increased phosphorylation upon reactivation. It is important to note that our prior work shows WT infection increases ERK phosphorylation relative to mock-infected myeloid cells; deletion of US28, however, further bolsters phospho-ERK, concomitant with robust viral replication, as US28 deletion virus fails to maintain latency (22, 25). Taken together with our findings herein, we propose a model whereby reactivation triggers an interaction (physical or proximity-mediated) between US28 and UL78 that renders US28 incapable of maintaining phospho-ERK suppression. Work aimed at deciphering the exact mechanism by which this occurs is underway.

Our data herein support prior transcriptional data showing UL78 is expressed during latency (**Fig. 1**) (11–14). Why then does UL78 expression only influence ERK activity upon reactivation and not during latency? Certainly, a complete understanding of other functions of latently-expressed UL78 remains outstanding, and it is indeed possible other functions will emerge. In line with our proposed model that UL78 regulates US28, which then alters signaling, we would further propose that US28’s other interactions during latency prevent the US28:UL78 crosstalk. During viral latency, US28 interacts with the cellular receptor tyrosine kinase, ephrin type-A receptor 2 (EphA2), which results in attenuated MAPK signaling to maintain latency (22). It is possible this interaction prevents UL78 from influencing US28 during latency. Alternatively, it is possible US28 and UL78 are not robustly expressed in the same location to a level above the threshold necessary for cross-talk, and by an unknown mechanism, reactivation results in their concentrated expression in close proximity, allowing UL78 to impact US28 function. Future work aimed at delineating these possibilities will further clarify this mechanism.

To date, UL78 remains a putative GPCR. UL78 has no known ligand(s) that bind this receptor and is thus considered an orphan receptor. Additionally, while many labs have interrogated the signaling functions of the CMV GPCRs for decades, UL78 has no known signaling functions to-date (27, 37), thus whether this is a functional GPCR remains elusive. It is, however, important to note that Medica, et al. presented findings at the 48^th^ Annual International Herpesvirus Workshop showing that UL78 couples Gα_i_ G proteins via the DRL motif in an overexpression system. Further, this group showed that mutation of the DRL motif to DAL rendered the virus unable to efficiently reactivate in CD34^+^ cell models (43). These differences could reflect a difference in CD14^+^ monocytes and less differentiated CD34^+^ cells. Alternatively, the tissue from which the material was derived (e.g., blood versus fetal liver) could also impact findings. Nonetheless, the requirement for UL78 during reactivation is a shared phenotype, thus highlighting the importance of this vGPCR to CMV biology.

Overall, our study unveils the first role of UL78 in myeloid cells and its regulation of cellular signaling in favor of CMV reactivation. This study established a foundation for future research aimed at more completely understanding the mechanisms by which UL78 influences latency and reactivation, thereby opening new avenues for therapeutic interventions to prevent viral reactivation.

## MATERIALS AND METHODS

### Cells and viruses

Newborn human foreskin fibroblasts (NuFF-1; GlobalStem, passage 13 - 25) and MRC-5 embryonic lung fibroblasts (ATCC; passage 10 – 27) were cultured in Dulbecco’s Modified Eagle’s Medium (DMEM), supplemented with 10% fetal bovine serum (FBS), 2 mM L-glutamine, 0.1 mM nonessential amino acids, 10 mM Hepes, and 100 U/ml each of penicillin and streptomycin. HEK 293T cells (ATCC) were cultured in DMEM, supplemented with 10% newborn calf serum (NCS), as well as 100 U/ml each of penicillin and streptomycin. THP-1 cells (ATCC) were maintained in RPMI 1640, supplemented with 10% FBS and 100 U/ml each of penicillin and streptomycin at a density of 5.0 ×10^5^ cells/ml. Primary CD14^+^ monocytes were isolated from de-identified cord blood samples (Abraham J. & Phyllis Katz Cord Blood Foundation d.b.a. Cleveland Cord Blood Center and Volunteer Donating Communities in Cleveland and Atlanta) via magnetic separation using the human CD14 MicroBeads (Miltenyi Biotec). Cells were cultured at 7.5 × 10^5^ cells/ml in RPMI 1640 (ATCC), supplemented with 100 U/ml each of penicillin and streptomycin and 1% heat-inactivated human serum (Millipore Sigma). All cells were maintained at 37°C and 5% CO_2_.

The bacterial artificial chromosome (BAC)-derived CMV isolate (TB40/E-BAC, clone 4) (45) was previously engineered to express mCherry (46). TB40/E*mCherry* (WT) was then used to generate a recombinant virus expressing a triple FLAG epitope tag in-frame with the UL78 ORF at its C-terminus, TB40/E*mCherry*-UL78-3xF, (UL78-3xF). Finally, UL78-3xF was used as the template to delete the entire UL78-3xF ORF, yielding TB40/E*mCherry*-UL78Δ (UL78Δ), both of which were previously characterized (28). UL78-3xF was also used as the template to generate two additional viruses by *GalK* recombineering (34, 35). Essentially, the galK gene was PCR amplified from pGalK using primers listed in **Table S1**. Recombination-competent SW105 *Escherichia coli* (*E. coli*) containing TB40/E*mCherry*-UL78-3xF were electroporated with the resulting product. The putative G protein-coupling domain was mutated at amino acid position 133-135 using a PCR amplified gBlock (**Table S1**), resulting in the change of the DRL motif to either AAA, DAL, or DRY). The resulting amplified gBlock was used to transform galK-positive clones, after which clones were counter-selected against galK, resulting in the virus mutants TB40/E*mCherry*-UL78^AAA^-3xF (UL78^AAA^), TB40/E*mCherry*-UL78^DAL^-3xF (UL78^DAL^), and TB40/E*mCherry*-UL78^DRY^-3xF (UL78^DRY^). The same process was applied to the C-terminus of the *UL78* gene in the TB40/E*mCherry*-US28-3xF backbone to introduce a triple HA epitope tag in-frame at the UL78 C-terminus, yielding TB40/E*mCherry*-US28-3xF:UL78-3xHA (US28-3xF:UL78-3xHA). All newly generated viruses were verified by Sanger sequencing using primers listed in **Table S1**.

### Virus propagation and growth assays

Virus was generated, propagated, and titered essentially as described elsewhere (34). In brief, BAC DNA was transfected into NuFF-1 cells, after which virus was expanded on naïve MRC-5 cells. Extracellular virus was collected at 100% cytopathic effect (CPE), concentrated by ultracentrifugation (71,800 x *g*, 90 minutes [min], 25°C) through a 20% sorbitol cushion (20% D-sorbitol, 1M Tris, pH 7.2, 1.0 M MgCl_2_), resuspended in X-VIVO 15 (Lonza) containing 1.5% BSA, flash frozen in liquid nitrogen, and stored at −80^°^C. Titers for viral stocks were quantified by tissue culture infectious dose 50 (TCID_50_) assay.

To quantify viral growth kinetics, NuFF-1 cells were infected at a multiplicity of infection (MOI) of 0.01 TCID_50_/cell. Cell-free virus was collected over a 16-day (d) time course. Additionally, cell-associated virus was collected at the final time point (16 d post-infection; dpi). All samples were stored at −80^°^C, and titers were determined by TCID_50_ assay.

### DNA and protein analyses

To isolate DNA, cells were lysed in TNE buffer (400 mM NaCl, 10 mM Tris pH 7.5, 10 mM EDTA), supplemented with 40 µg proteinase K (ThermoFisher) and 6.4 µg sodium dodecyl sulfate (SDS; VWR), then mixed by vortexing, and incubated overnight at 37^°^C. DNA was then extracted using phenol/chloroform (Fisher), by adding equal volume to each sample and centrifuged at 14,000 x *g* for 5 min at 4°C. Afterwards, 40 µg RNaseA (Roche) was added and incubated for 1 hour (h) at 37°C, and then extracted with equal volume phenol/chloroform (Fisher) and equal volume chloroform (Invitrogen), as above. DNA was precipitated at −20°C by adding equal volume of 100% ice cold ethanol to each sample tubes for 1 h, followed by centrifugation at 14,000 x *g* for 30 min at 4°C. Resulting pellets were washed with 70% ethanol and centrifuged at 21,300 x *g* for 5 min at 4°C. DNA pellets were air dried at room temperature for 10 min and resuspended in 10 mM Tris, pH 8.0. Viral and cellular DNA was quantified by qPCR using primers directed at UL69 and MDM2, respectively (**Table S2**). Genome copy numbers were extrapolated from a standard curve generated from serial dilutions of known quantities of a BAC-standard, which also contains MDM2 sequence (21). Samples were analyzed in triplicate using a 96-well format CFX Connect Real-Time PCR machine (Bio-Rad).

For protein analyses, cells were lysed in RIPA buffer (1% NP-40, 1% sodium deoxycholate, 0.1% SDS, 0.15 M NaCl, 0.01 M NaPO_4_, 2.0 mM EDTA, pH8.0) adjusted to a final pH of 7.2 with NaOH, supplemented with protease (Merk) and phosphatase (Roche) inhibitor cocktails for 1 h on ice, vortexing every 15 min. Protein concentration was determined by Bradford assay using Protein Assay Reagent Concentrate (Bio-Rad), according to the manufacturer’s instructions. Samples used to detect vGPCRs were denatured at 42°C for 10 min; all other samples were denatured at 95^°^C for 10 min. Protein samples were separated by SDS-PAGE, transferred to nitrocellulose membrane (Cytiva) by semi-dry transfer, and detected using the following antibodies: anti-FLAG, clone M2 (Millipore Sigma, 1:7,500); anti-HA, clone 29F4 (Cell Signaling Technology (CST)); anti-V5, V8137-2MG (Sigma, 1:2,000); anti-phospho-p44/42 MAPK (ERK1/2) (CST, 1:1,000); anti-p44/42 MAPK (ERK1/2) (CST, 1:1,000); anti-β-actin peroxidase (MilliporeSigma, 1:20,000); goat anti-rabbit and goat anti-mouse HRP secondary antibodies (Jackson Immuno-Research Labs, 1:10,000).

### Latency and reactivation assays

Primary CD14^+^ monocyte cells were isolated from umbilical cord blood by magnetic separation using human CD14^+^ Microbeads (Miltenyi Biotec), according to the manufacturer’s instructions. Cells were infected (MOI = 1.0 TCID_50_/cell) by centrifugal enhancement (1,000 x *g*, 30 min, room temperature) and incubated overnight at 37°C and 5% CO_2_. Next, cells were washed three times with 1X PBS and cultured at 3 × 10^6^ cells/ml in X-VIVO 15. At 7 dpi, half of each infected cell population was serially diluted in triplicate in reactivation media (RPMI 1640, 10% FBS, 10 ng/ml macrophage colony stimulating factor [MCSF]), or maintained in latency media (X-VIVO 15). Cells were then co-cultured on NuFF-1 cells and were maintained for 14 d. The frequency of infectious centers was quantified by extreme limiting dilution analysis (ELDA); https://bioinf.wehi.edu.au/software/elda/ (36), essentially as previously described (47).

THP-1 cells were infected (MOI = 1.0 TCID_50_/cell) by centrifugal enhancement (1,000 x *g*, 30 min, room temperature) in low serum media (X-VIVO 15), then incubated for 90 min at 37°C with 5% CO_2_. Virus inocula was then removed, and cells washed three times with 1X PBS, replenished with X-VIVO 15, and returned to culture as indicated in the text.

### Co-immunoprecipitation (co-IP) assays

NuFF-1 cells were infected (MOI = 0.5 TCID_50_/cell) with TB40/E*mCherry*-US28-3xF:UL78-3xHA (US28-3xF:UL78-3xHA) for 96 h, and were then lysed on ice for 2 h with IP buffer (150 mM NaCl, 25 mM Tris, pH 7.4, 10 mM MgCl, 2 mM EDTA, 1% Triton X-100), supplemented with complete protease inhibitor cocktail (Merk). Cells were centrifuged at 12,000 x *g* for 10 min at 4°C, after which a 10% aliquot was reserved as an input control. Indicated proteins were precipitated overnight on a nutating rack with equal volume of antibody (anti-HA, CST; rabbit IgG, MilliporeSigma; mouse IgG, CST) and protein G Sepharose beads (Sigma Aldrich) at 4°C. Beads were then washed four times at 4°C in IP wash buffer, and 10% of the first and fourth washes were retained for analyses. All samples were denatured at 42°C for 10 min and immunoblotted, as above.

### Immunofluorescence assays (IFAs)

NuFF-1 cells were grown on 22 x 22 mm coverslips (Fisher) overnight, and infected (MOI = 0.5 TCID_50_/cell) as indicated in the text. For IFAs of THP-1 cells, cells were infected (MOI = 1.0 TCID_50_/cell) by centrifugal enhancement (1,000 x *g*, 30 min, room temperature). At 5 dpi, cells were cushioned onto Ficoll-Plaque PLUS (Cytiva) at 450 x *g* for 35 min (no brake), washed three times with 1X PBS, replenished with X-VIVO 15, and cultured in 6-well plates containing 22 x 22 mm coverslips (Fisher). At 7 dpi, cells were treated for an additional 2 d with 20 nM 12-Otetradecanoylphorbol-13-acetate (TPA) to induce cellular differentiation or vehicle (DMSO; v/v) to maintain latent conditions. Alternatively, THP-1 cells were pre-differentiated with 20 nM TPA for 24 h on coverslips to induce cellular differentiation. Cells were then washed with 1X PBS, replenished with X-VIVO 15, and infected as above for 48 h as indicated in the text. In each case, cells were processed as above for IFA.

Regardless of cell type, cells were fixed in 2% paraformaldehyde at 37°C, permeabilized with 0.25% saponin at 25°C, and blocked with 10% human serum and 0.25% saponin in 1X PBS. Cells were stained with the following antibodies: anti-HA (CST, 1:1,000); anti-FLAG (MilliporeSigma, 1:1,000); Alexa 488-conjugated anti-mouse (Abcam, 1:1,000), Alexa 647-conjugated anti-rabbit (Abcam, 1:1,000). Nuclei were visualized with 4’-6’-diamidino-2-phenylindole (DAPI; Fisher; 1:5,000). Coverslips were mounted with FlourSave Antifade reagent (ThermoFisher), and images were captured using a Leica TCS-SP8-AOBS inverted confocal microscope and Leica Application Suite X (LAS X) Software, version 3.5.5.

Where indicated, colocalization of viral proteins was performed digitally using ImageJ/Fiji software plugin coloc2 (version 1.54p) (48). Colocalization was quantified using the pixel intensity correlation over space method based on Pearson’s correlation coefficient across the number of view fields indicated in the text.

### Generation and expression of vGPCR-expressing plasmids

The expression vector backbones, pCMS(crs-)-eGFP and pCMS(crs-)-dsRed, were kind gifts from Eain Murphy (SUNY Upstate Medical University). pCMS(crs-)-eGFP-US28-3xF contains the US28 ORF with an in-frame triple FLAG (3xF) epitope tag on the C-terminus. pCMS(crs-)-dsRed-UL78-V5 contains the UL78 ORF with a single V5 epitope tag on the C-terminus. The primers used to generate these constructs, as well as those used to verify their sequences, are listed in **Table S2**.

MRC-5 cells were transfected with vGPCR-containing pCMS vectors using the lipofectamine 2000 kit (Invitrogen), following the manufacturer’s instructions. Cell lysates were collected 48 h post-transfection, and the expression of US28 and UL78 were assessed by immunoblot for their respective tags.

### Statistical analyses

Data were analyzed using GraphPad Prism (version 10.1.2) software (GraphPad Software LLC) and Excel (Microsoft). The statistical test performed for each experiment is indicated in the text.

## ACKNOWLEDGEMENTS

This work was supported by the National Institutes of Health (R01AI153348 to C.M.O’C.). The funders had no role in study design, data collection and interpretation, or the decision to submit the work for publication.

## Notes

### Competing Interest Statement

The authors have declared no competing interest.

