## Supplemental Figures for "Human cytomegalovirus-encoded G protein-coupled receptor (GPCR), UL78, regulates viral reactivation"

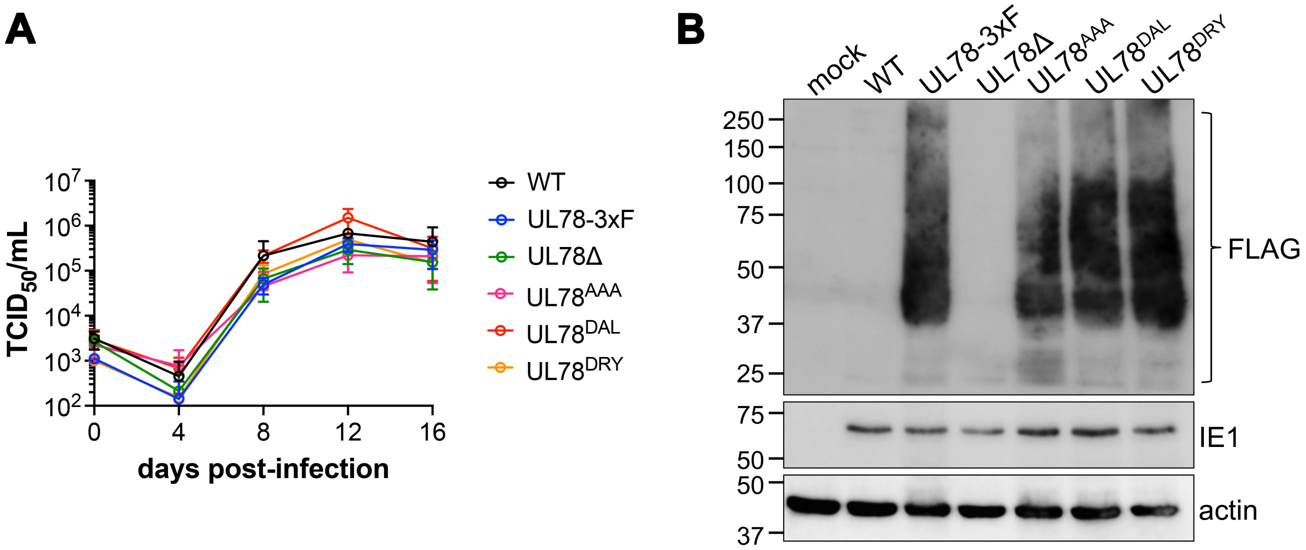


**Figure S1. Recombinant and deletion UL78 viruses display wild type lytic replication kinetics in fibroblasts. (A)** Fibroblasts (MRC-5) were infected with the indicated viruses, supernatants collected over a 16-day time course and then analyzed by $TCID_{50}$ assay. N=3 biological replicates, each with 3 technical replicates for each virus at each time point. Error bars represent SD of the mean; significance calculated using t-test. Data is not significant. **(B)** Fibroblasts (MRC-5) were mock-, WT-, UL78-3xF, UL78Δ-, $UL78^{AAA}$-, $UL78^{DAL}$-, or $UL78^{DRY}$-infected (MOI = 0.5 $TCID_{50}$/cell). Lysates were collected 96 hpi and probed for FLAG (to detect UL78), IE1, or actin (loading control). N=3, representative blots shown.

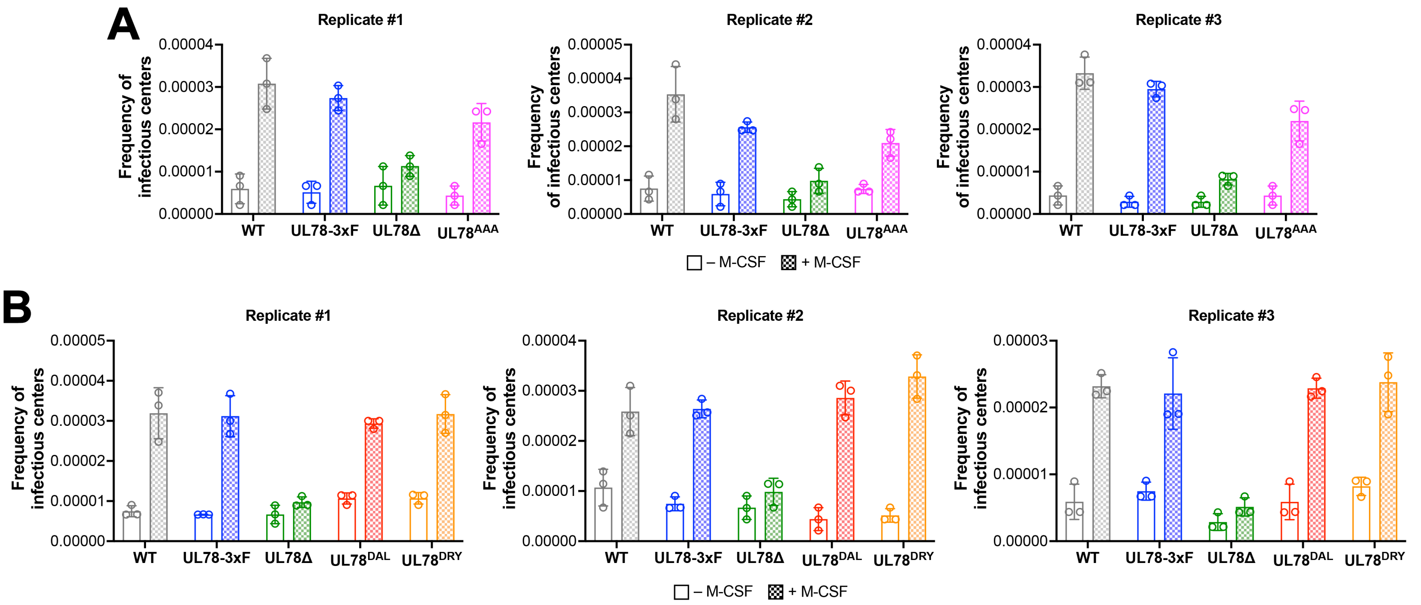


**Figure S2. Each independent replicate for combined ELDA data in Figure 2. (A)** Data corresponds to Fig. 2A. **(B)** Data corresponds to Fig. 2B. **(A,B)** Primary CD14$^+$ cells were infected with the indicated viruses at MOI = 1.0 TCID$_{50}$/cell and cultured for 7 d in conditions favoring latency. CD14$^+$ cells were maintained under latent conditions (- M-CSF) or treated with M-CSF (+ M-CSF) to differentiate the cells. Cells were then co-cultured with naïve fibroblasts, and following 14 d in co-culture, ELDA was used to quantify reactivation. Data points (open circles) represent technical replicates. Error bars represent SD of the mean.

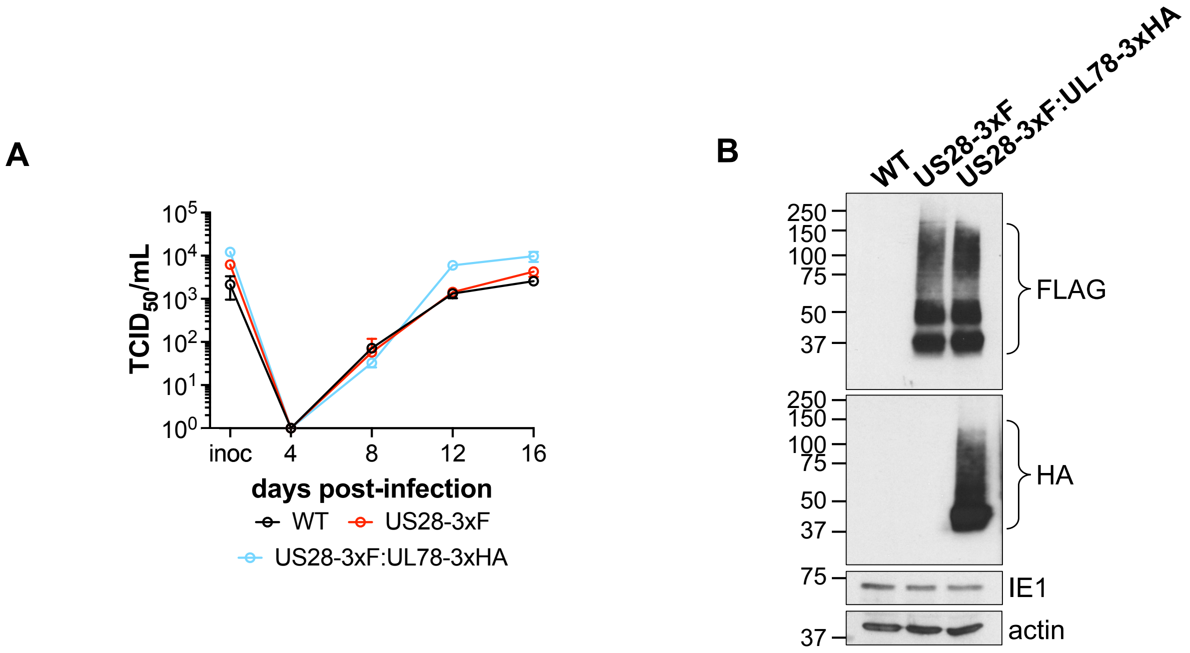


**Figure S3. TB40/E*mCherry*-US28-3xF:UL78-3xHA (US28-3xF:UL78-3xHA) infection of fibroblasts displays wild type growth kinetics. (A)** Fibroblasts (NuFF-1) were infected with the indicated viruses (MOI = 0.01 $TCID_{50}$/cell). Supernatants were collected at the indicated time points and titers assessed by $TCID_{50}$ assay. N=2 biological replicates; each with three technical replicates per condition. Representative assay shown. Error bars represent SD of the mean. **(B)** Fibroblasts (NuFF-1) were infected with the indicated viruses (MOI = 0.5 $TCID_{50}$/cell) for 96 h. Lysates were collected and probed for FLAG (to detect US28), HA (to detect UL78), IE1, and actin. N=2; representative blots shown.

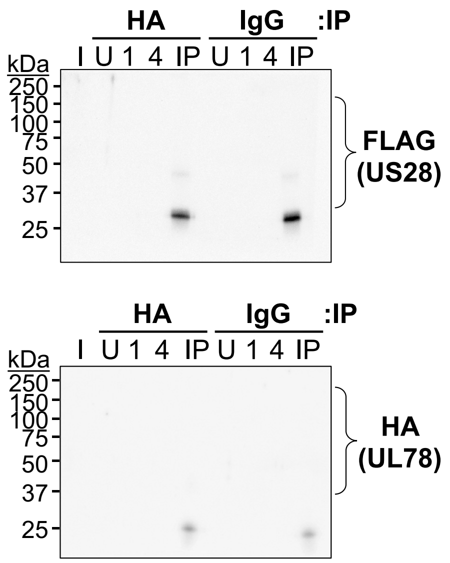


**Figure S4. TB40/E*mCherry* control data for Figure 4.** NuFF-1 fibroblasts were infected (MOI = 0.5 $TCID_{50}$/cell) with TB40/E*mCherry.* Cells were harvested 96 hpi and lysates were immunoprecipitated (IP) with either anti-HA or anti-IgG antibodies and immunoblotted for (**top**) FLAG to detect US28 or (**bottom**) HA to detect UL78**.** I, input; U, unbound; 1, 1st wash; 4, 4th wash; IP, immunoprecipitated sample. Representative blots shown.

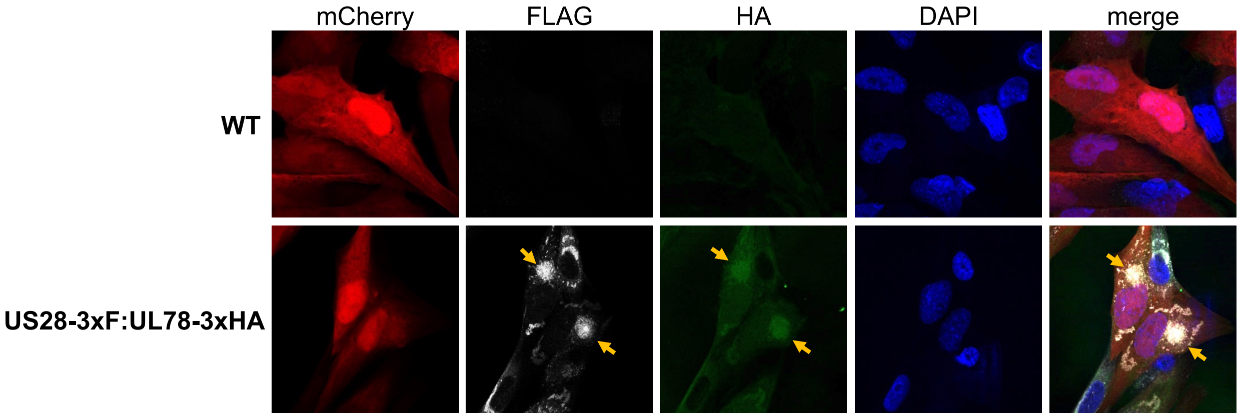


**Figure S5. UL78 and US28 colocalize in the VAC in lytically infected fibroblasts.** NuFF-1 fibroblasts were infected with either TB40/E*mCherry* (wild type, WT) or TB40/E*mCherry*-US28-3xF:UL78-3xHA (MOI = 0.5 $TCID_{50}$/cell) for 96 h. Cells were fixed, permeabilized, and then stained for US28 with antibodies directed at the triple FLAG epitope (white) and UL78 with antibodies directed at the triple HA epitope (green). mCherry (red) serves as a marker of infection and DAPI (blue) was used to visualize cell nuclei. Gold arrows denote areas of US28 and UL78 co-localization. Images were acquired using a 63x objective. N= 2; representative images shown.

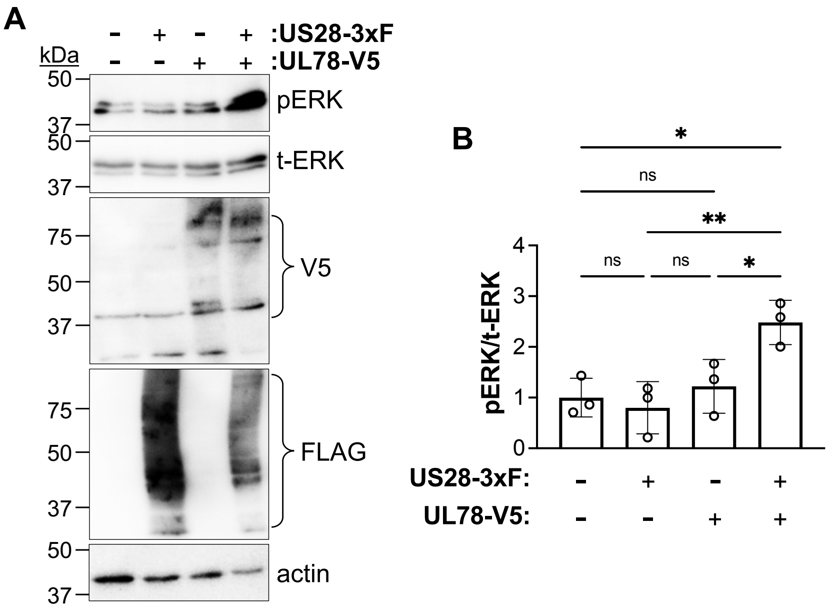


**Figure S6. Co-expression of UL78 and US28 alters US28-mediated signaling. (A)** Fibroblasts (MRC-5) were transfected with US28-3xF or UL78-V5 expressing constructs alone or in combination. Cells were collected at 2 d post-transfection, and lysates were probed with antibodies directed at phosphorylated ERK (p-ERK), total ERK (t-ERK), V5 (to detect UL78), FLAG (to detect US28), or actin. N = 3; representative blots shown. **(B)** Data from (**A**) were analyzed for the pERK to t-ERK phospho-ratio. Each data point represents densitometry values from each of the 3 biological replicates. Data is shown relative to the pERK/t-ERK ratio in control cells. Error bars are SD of the mean; significance was calculated using one-way ANOVA.

**Table S1. Oligonucleotides used to generate viral recombinants.**

| Primer use | Sequence* | Target** |
|---|---|---|
| Generating galK insertion (for UL78$^{AAA}$, UL78$^{DAL}$, UL78$^{DRY}$) | CGACGTGGGCCTATATTCGACGGCGTTGTTTTTCCTCTTTCTGATACTGGCCTGTTGACAATTAATCATCGGCA | UL78 galK For |
| | GTTTTCGCGCGTCTCATGATGCCAGAGATCACGGCCGTAAGATATGGCCGTCAGCACTGTCCTGCTCCTT | UL78 galK Rev |
| UL78$^{AAA}$ insert | TCTTTGTCGACGACGTGGGCCTATATTCGACGGCGTTGTTTTTCCTCTTTCTGATACTGGCAGCTGCGTCGGCCATATCTTACGGCCGTGATCTCTGGCATCATGAGACGCGCGAAAACGCCGGCGTGGCG | UL78$^{AAA}$ gBlock |
| UL78$^{DAL}$ insert | TCTTTGTCGACGACGTGGGCCTATATTCGACGGCGTTGTTTTTCCTCTTTCTGATACTGGATGCACTGTCGGCCATATCTTACGGCCGTGATCTCTGGCATCATGAGACGCGCGAAAACGCCGGCGTGGCG | UL78$^{DAL}$ gBlock |
| UL78$^{DRY}$ insert | TCTTTGTCGACGACGTGGGCCTATATTCGACGGCGTTGTTTTTCCTCTTTCTGATACTGGATCGTTACTCGGCCATATCTTACGGCCGTGATCTCTGGCATCATGAGACGCGCGAAAACGCCGGCGTGGCG | UL78$^{DRY}$ gBlock |
| UL78 gBlock amplification primers | TCTTTGTCGACGACGTGGGC | UL78 gBlock For |
| | CGCCACGCCGGCGTTTTCGC | UL78 gBlock Rev |
| Generating galK insertion (for UL78-3xHA) | GCACCGACGGCGAAAACACCGTCGCGTCCGACGCAACGGTGACGGCATTACCTGTTGACAATTAATCATCGGCA | UL78-3xHA galK For |
| | GTGATTTATCTGCCACTTTTCTCCCCGCTGCCGTACAGCGCCGCCGCTCATCAGCACTGTCCTGCTCCTT | UL78-3xHA galK Rev |
| UL78-3xHA insert | GCACCGACGGCGAAAACACCGTCGCGTCCGACGCAACGGTGACGGCATTATACCCATATGACGTTCCAGACTACGCGTATCCGTACGACGTTCCGGATTACGCTTACCCTTACGACGTACCTGACTACGCTTGAGCGGCGGCGCTGTACGGCAGCGGGGAGAAAAGTGGCAGATAAATCAC | UL78-3xHA gBlock |
| UL78-3xHA gBlock amplification primers | TGCACCGACGGCGAAAACACCG | UL78-3xHA gBlock For |
| | GTGATTTATCTGCCACTTTTCTCC | UL78-3xHA gBlock Rev |
| Sequencing primers UL78 | GCTGCATAAAGGGCGAACG | UL78 seq For |
| | TGGCTAACGACGTGTGAACC | UL78 seq Rev |

*Primer sequences shown the 5' to 3' orientation. Underlined sequences correspond to the pGalK plasmid. **For, forward; Rev, reverse.

**Table S2. Oligonucleotide pairs used for PCR.**

| Target | Forward (5’- 3’)* | Reverse (5’-3’)* |
|---|---|---|
| UL69 | GGCTGATGATCTTGCGGGAA | CGAGAGTCTACGTCTGGCAC |
| MDM2 | CCCCTTCCATCACATTGCA | ACCCACTCCTCCACCTTTGAC |
| US28 ORF | GGGCTCGAGATGACACCGACGACGACG | CCCCTCTAGATTACTTGTCGTCGTCGTCCTTGTAGTCGAACCCCTTGTCGTCGTCGTCCTTGTA |
| UL78 ORF | GCCGCTCGAGATGGGTAAGCCTATCCCTAACCCTCTCCTCGGTCTCGATTCTACGTCCCCTTCTGTGGAGGAGACTAC | CGACTCTAGATCACGTAGAATCGAGACCGAGGAG |

*Underlined sequences correspond to restriction sites in the primers: forward, XhoI and reverse, XbaI.
